# Acute inflammation is a predisposing factor for weight gain and insulin resistance

**DOI:** 10.1101/583773

**Authors:** Edson M. de Oliveira, Jacqueline C. Silva, Thais P. Ascar, Silvana Sandri, Alexandre F. Marchi, Silene Migliorini, Helder T. I. Nakaya, Ricardo A. Fock, Ana Campa

## Abstract

**Aim:** Intense endotoxaemia and infection are able to reduce appetite and induce a catabolic state, therefore leading to weight loss. However, it is underexplored its late effects on energy homeostasis, regulation of body weight and glucose metabolism. Here we addressed whether serial intense endotoxaemia, characterized by an acute phase response and weight loss, could be an aggravating or predisposing factor to diet-induced obesity (DIO) and associated metabolic impairments.

**Methods:** Male Swiss Webster mice were submitted to 8 consecutive doses of lipopolysaccharide (LPS - 10 mg/kg), followed by 10 weeks in high-fat diet (HFD).

**Results:** After the end of the acute endotoxaemia period, mice under chow diet recovered their weight rapidly, within one-week recovery period, which remained similar to its control counterparts. However, acute endotoxaemia caused a long-lasting adipose tissue expression of the inflammatory markers TLR-4, CD14 and serum amyloid A (SAA) and, when challenged by a HFD, LPS-treated mice gained more weight, showed increased fat depots, leptin and insulin levels, and also impaired insulin sensitivity.

**Conclusions:** LPS-treated mice showed a higher susceptibility to the harmful effects of a subsequent HFD. Conditions leading to intense and recurrent endotoxaemia, such as common childhood bacterial infections, may resound for a long time and aggravate the effects of a western diet. If confirmed in humans, infections should be considered an additional factor contributing to obesity and type 2 diabetes epidemics and additionally impose more rigorous dietary recommendations for patients in post-infection recovery.

**Bullet points:**

- Intense endotoxemia causes a long-lasting increase in the expression of inflammatory markers in adipose tissue.
- Intense endotoxemia is a predisposing factor to diet-induced obesity and insulin resistance.
- Infections may contribute to weight gain when associated to a western diet.

## 1. Introduction

Metabolic endotoxaemia is the low-grade endotoxaemia derived from intestinal microbiota and modulated by high-fat feeding (1,2). Since long-term endotoxaemia has a role in weight gain and insulin resistance, we hypothesized that a transient and intense endotoxaemia also impacts on adipose tissue and weight gain.

Intense endotoxaemia is observed along bacterial infections and it is experienced multiple times during human life. It is characterized by a large catabolic process that frequently leads to weight loss (3). In mice, endotoxaemia mimicking bacterial infections may be achieved by endovenous or intraperitoneal administration of high doses of lipopolysaccharide (LPS), resulting in a pronounced inflammatory response characterized by a huge increment in serum levels of acute phase proteins, such as serum amyloid A (SAA) and C-reactive protein (CRP). Acute phase proteins are commonly used in clinical practice as an unspecific serum marker to confirm an inflammatory acute phase response. For instance, the serum levels of SAA may increase up to 1000-fold compared to a non-inflammatory state (4). Although in this study SAA was used as an inflammatory marker, it is mandatory to consider that SAA affects adipose tissue biology (5) and may be considered as a trigger driving obesity and insulin resistance, as we recently showed using SAA-targeted antisense oligonucleotide in an experimental model of diet-induced obesity (6).

Although it is known that intense endotoxaemia is promptly associated with weight loss, here we wonder if acute endotoxaemia was able to prime the adipose tissue favouring a more pronounced response to an obesogenic stimulus. To reach our goal, we used an experimental model of multiple inductions of acute endotoxaemia (consecutive LPS challenges) followed by a HFD period. Also, in order to verify if acute infection could be correlated with cell proliferation, adipogenesis and inflammation in adipose tissue, therefore predisposing it for future hypertrophy, we performed a Gene Set Enrichment Analysis using a publicly database relative to the effect of gram-negative bacteria infections on adipose tissue.

If intense endotoxaemia leads to biochemical changes involved in adipose tissue hyperplasia and hypertrophy, acute inflammation should be recognized as an aggravating factor for weight gain and obesity comorbidities.

## 2. Material and methods

### 2.1 Animals

Male Swiss Webster mice (21 days of age) were obtained from the Animal Facility of the Faculty of Pharmaceutical Sciences, University of São Paulo, Brazil, under approval by its Ethical Committee (CEEA n°297). The animals were housed inside standard polypropylene cages in a room maintained at 22±2°C in 12:12 h light/dark cycle (lights on at 7:00 am and off at 7:00 pm) and a relative humidity of 55±10%. Body weight was measured once a week during the entire protocol. Food and water intake were kept *ad libitum* and were measured every 2 days. Euthanasia occurred by anaesthesia overdose and ensured by cervical dislocation.

### 2.2 Acute endotoxaemia followed by recovery period under chow diet

The method of multiple inductions of acute endotoxaemia comprises intraperitoneal administration of 8 consecutive injections (every 3 days) of LPS 10 mg/kg (Lipopolysaccharides from *Escherichia coli* 026:B6, Sigma-Aldrich^®^, St. Louis, MO, USA), in saline (NaCl 0,9%), starting at weaning (21 days of age) with end at 45 days of age of the animal. The time between two acute endotoxaemia (3 days) was defined by the SAA production profile. It was observed that after LPS injection, SAA concentration increases over a 100-times, with maximum values in 12 hours, approximately 1500 μg/mL, with return to basal in 72 hours (Supplementary Table 1). For acute endotoxaemia experiments, mice were randomly assigned into 2 different groups: the Control and the LPS groups, with euthanasia occurring after the last acute phase period or after 6 weeks from the last acute phase induction (recovery period). The experimental design is illustrated in Supplementary Fig. 1a.

### 2.3 Acute endotoxaemia followed by High-Fat Diet (HFD)

For acute endotoxaemia followed by 10 weeks on a high-fat diet (LPS+HFD) experiments, the animals were randomly assigned into 2 different groups: HFD group and LPS+HFD group. The HFD mice were submitted to a HFD for 10 weeks starting concurrently with the LPS+HFD group. The LPS+HFD mice were underwent to multiple inductions of acute endotoxaemia followed by 1 week of recovery period in standard chow diet plus 10 weeks on a 30% HFD. In our experimental model, we consider the recovery period as 7 days, been the time that we observed weight reestablishment. The diet was produced following the American Institute of Nutrition’s recommendations for the adult rodent and its composition is listed in Supplementary Table 2. Body weight was measured every 3 days during acute phase period. The experimental design is illustrated in Supplementary Fig. 1b.

### 2.4 Glucose and insulin tolerance tests and measurements of serum leptin, adiponectin, insulin, IGF-I, SAA and endotoxin

i.p. Glucose and insulin tolerance tests (IPGTT and IPITT) were performed as described previously (7). Serum concentrations of the proteins below were determined using ELISA following the manufacturer’s instructions: leptin, adiponectin and insulin (Millipore^®^ Corporation, Billerica, MA, USA), SAA (Tridelta Development Ltd, Maynooth, Ireland) and IGF-I (R&D Systems^®^, Minneapolis, MN, USA). Endotoxin was measured with the Limulus Amoebocyte Lysate (LAL) chromogenic end-point assay (Lonza^®^, Allendale, NJ, USA).

### 2.5 Histological Analysis

Paraffin-embedded sections (5 μm thick) from epididymal adipose tissue were stained by haematoxylin and eosin to assess morphology. Immunohistochemistry for F4/80^+^ was performed using a rat antimouse F4/80^+^ antibody (1:100 dilution, AbD Serotec^®^, Raleigh, NC, USA) subsequently incubated with the appropriate secondary biotinylated antibody (Vector Laboratories Inc., Burlingame, CA, USA) and visualized with Immpact AEC peroxidase (Vector Laboratories Inc., Burlingame, CA, USA). Immunofluorescence for F4/80^+^, SAA and perilipin were performed using a rat anti-mouse F4/80^+^ antibody and rabbit anti-mouse perilipin (both 1:100 dilution, Abcam^®^, Cambridge, UK), and a rabbit anti-mouse SAA (1:200 dilution, kindly provided by Dr. de Beer laboratory, University of Kentucky, KY, USA), subsequently incubated with the appropriate secondary fluorescent antibody (Invitrogen^®^, Camarillo, CA, USA) and the slides mounted using Vectashield set mounting medium with 4,6-diamidino-2-phenylindol-2-HCl (DAPI; Vector Laboratories Inc., Burlingame, CA, USA). An isotype control was used to ensure antibody specificity in each staining. Tissue sections were observed with a Nikon Eclipse 80i microscope (Nikon^®^) and digital images were captured with NIS-Element AR software (Nikon^®^).

### 2.6 In vivo peripheral fat area quantification

Two X-ray images were taken at different energy levels allowing circumscribe the adipose tissue on the animals as described previously (7).

### 2.7 Quantitative Real-Time PCR

Total RNA from epididymal adipose tissue was isolated using Qiagen RNeasy^®^ Lipid Tissue Mini kit (Qiagen, Hilden, Germany). cDNA was then synthesized from 1 μg of RNA using the High Capacity cDNA Reverse Transcription (Life Technologies^®^, Grand Island, NY, USA). Real-time PCR were performed using SyBr^®^ Green Master Mix (Life Technologies^®^, Grand Island, NY, USA). The primer sequences are detailed in Supplementary Table 3. Real-time PCR for *Saa3* was performed using the TaqMan^®^ assay (Applied Biosystems^®^, Grand Island, NJ, USA), catalogue number Mm00441203_m1 – *Saa3* and β-actin (*Actb*), number 4552933E, as an endogenous housekeeping gene control. Relative gene expression was determined using the 2^-ΔΔCt^ method.

### 2.8 Gene Set Enrichment Analysis of publicly available microarray data

We collected from GEO (http://www.ncbi.nlm.nih.gov/geo, GSE50647) the expression profiles of mouse visceral adipose tissue. In the study (8), authors exposed chow-fed apolipoprotein E (apoE) deficient mice to either 1) recurrent intravenous infection with *A. actinomycetemcomitans* or 2) a combination of recurrent intravenous infection with *A. actinomycetemcomitans* with a chronic intranasal infection with *C. pneumonia*. For the Gene Set Enrichment Analysis (GSEA) we ranked genes based on their mean log_2_ fold-change values between infected compared to uninfected mice. We then utilized custom gene sets, which contained genes related to: proliferation, adipogenesis, inflammation and SAA family. GSEA was performed using default parameters. Heat maps were used to display all genes from a statistically significant gene set.

### 2.9 Statistical analysis

Results were presented as mean ± SEM. Statistical analysis was performed with Graph Pad Prism4 (Graph Pad Software, Inc., San Diego, CA, USA). Comparisons between two groups were conducted with the unpaired Student’s *t* test. Data with two independent variables were tested by two-way analysis of variance with Bonferroni *post hoc* test. The level of significance was set at \**p* < 0.05.

## 3. Results

### 3.1 Acute endotoxaemia affects adipose tissue but does not lead to mice weight gain under chow diet

In order to verify the effect of acute endotoxaemia in adipose tissue, mice were subjected to 8 consecutives LPS challenges, every 3 days (Supplementary Fig. 1a). During the acute phase response, endotoxin (Fig. 1a) and SAA (Fig. 1b) levels raised over a hundredfold in serum and mice developed overt signs of endotoxaemia (hunched posture, reduced movement and piloerection) with no animal death. It is known that acute endotoxaemia change food intake behaviour causing a reduced food intake and leading to weight loss. During the acute endotoxaemia period, LPS animals showed a reduced caloric intake (up to 40% lower; Fig. 1c), causing weight loss (approx. 12.5% of their total weight; Fig. 1d) and 20% of epididymal adipose tissue mass (Fig. 1e).

**Figure 1.**
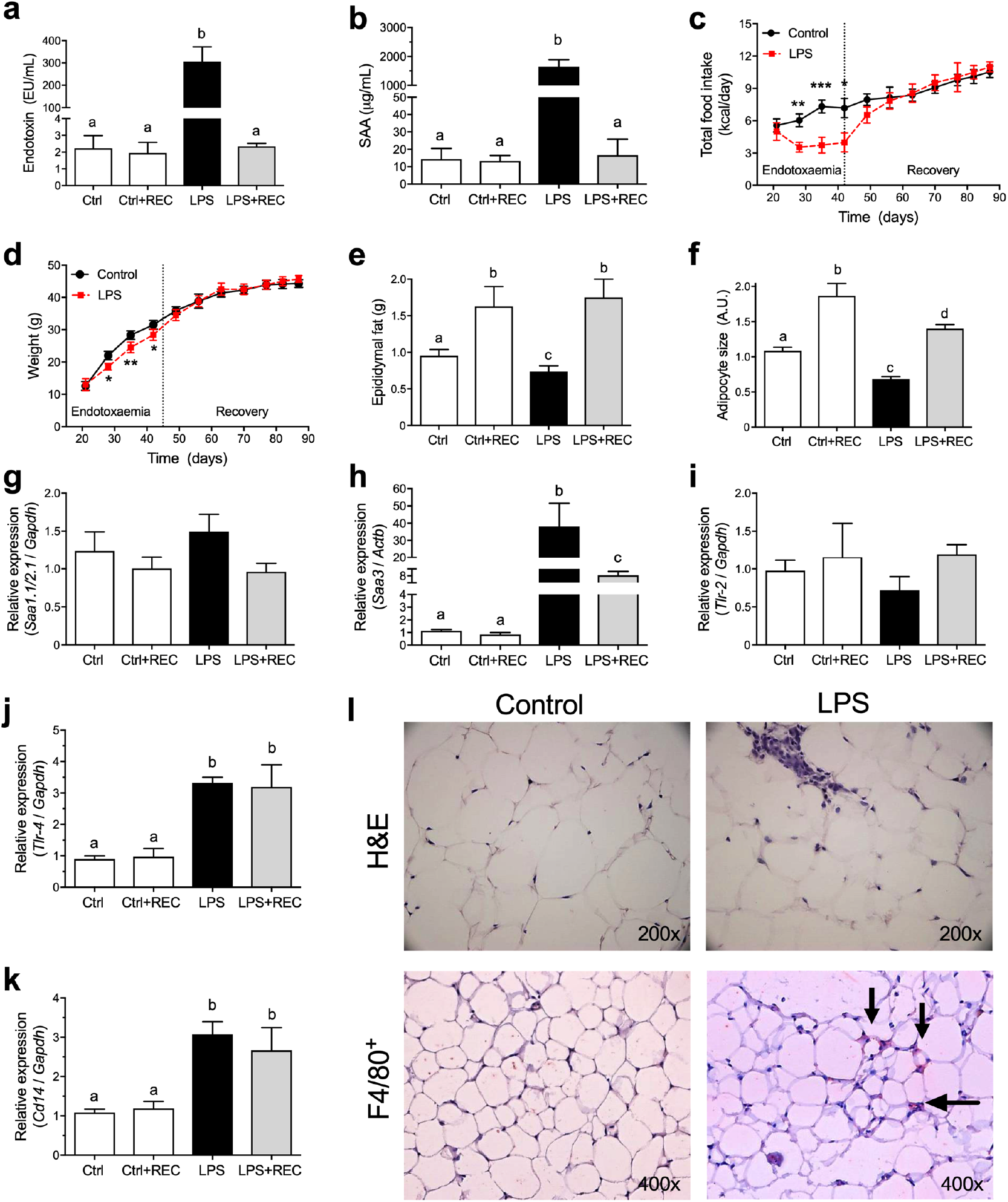
Acute endotoxaemia affects adipose tissue but does not lead to mice weight gain under chow diet. Swiss Webster mice were submitted to i.p. administration of 8 consecutive doses of 10 mg/kg LPS, every 3 days. **(a)** Endotoxin and **(b)** SAA concentration in serum. **(c)** Daily caloric intake. **(d)** Weight gain curve of Control and LPS mice. The vertical dashed line in *c* and *d* indicate the end of acute endotoxaemia period. **(e)** Epididymal fat pad weight. **(f)** Epididymal adipocyte size. **(g-k)** Quantitative Real-Time PCR was performed to assess mRNA expression of **(g)** *Saa1.1/2.1*, **(h)** *Saa3*, **(i)** *Tlr-2*, **(j)** *Tlr-4* and **(k)** *Cd14* in epididymal adipose tissue. **(l)** Histological sections of epididymal fat pads after LPS challenges showing adipocyte morphology on haematoxylin and eosin staining and macrophage infiltration (F4/80^+^). Data are means ± SEM from 6 mice per group (\**p* < 0.05, \*\**p* < 0.01, \*\*\**p* < 0.001, between groups, as indicated).

By the histological analysis from epididymal adipose tissue (Fig. 1l), it was also possible to verify a decrease in 30% of the adipocyte size in the LPS group (Fig. 1f). Besides weight loss, mice had increased inflammatory markers, such as macrophage infiltration (F4/80^+^ cells, Fig. 1l) and *Saa3* mRNA expression (Fig. 1h) without *Saa1.1* and *Saa2.1* modulation in adipose tissue (Fig. 1g). The expression of *Tlr-4* (Fig. 1j) and *Cd14* (Fig. 1k) had an increase of more than three times, although the *Tlr-2* (Fig. 1i) remained unaltered.

During the experimental protocol, no tolerance to LPS was observed, with increment in serum endotoxin (Supplementary Figure 2a) and SAA (Supplementary Figure 2b) remaining similar after each LPS challenge. After the endotoxaemia period, the animals recovered their weight in the course of a week without showing any difference from Control group in the 42 consecutive days (6 weeks) (Fig. 1c and d). One week after the last LPS challenge (LPS + REC) was also sufficient for the reestablishment of serum levels of endotoxin (Fig 1a) and SAA (Fig 1b), however, the expression of *Saa3* (Fig 1h), *Tlr-4* (Fig 1j) and *Cd14* (Fig 1k) remained augmented in the adipose tissue.

### 3.2 A previous history of acute endotoxaemia exacerbates HFD complications

After the LPS challenges and a recovery period of one week, mice were submitted to a HFD for 70 days (10 weeks) (Supplementary Fig. 1b). HFD-fed mice without having previously been submitted to LPS challenges were used as controls. The shift of chow diet to HFD resulted in an increment of approximately twice in the caloric intake for both groups, HFD and LPS+HFD (Fig. 2a). Despite the similar caloric intake between these groups, mice previously submitted to multiple LPS challenges (LPS+HFD) showed a different growth curve with increased total body weight (approximately in 15%) (Fig. 2b), with increased epididymal (Fig. 2c) and subcutaneous (Fig. 3e) adipose tissue depots, and no difference in retroperitoneal (Fig. 3d) fat. The data were confirmed using X-rays images highlighting the subcutaneous fat area on the animals, showing that LPS+HDF mice have a higher peripheral fat area, an increment in 23% in adipose tissue (Fig. 2f and g). Besides that, the epididymal fat from LPS+HFD mice presented larger adipocytes than HFD group (Fig. 2h), that may explain the metabolic phenotype. On the other hand, there is no difference in the adipose tissue inflammatory profile, showing similar macrophage infiltration (F4/80^+^) and SAA production (Fig. 2i) in both HFD and LPS+HFD groups. However, it is important to highlight that all aforementioned parameters (adipocyte hypertrophy, macrophage infiltration and SAA production in the adipose tissue) are increased when compared to lean control group (under chow diet) (Fig. 2i).

**Figure 2.**
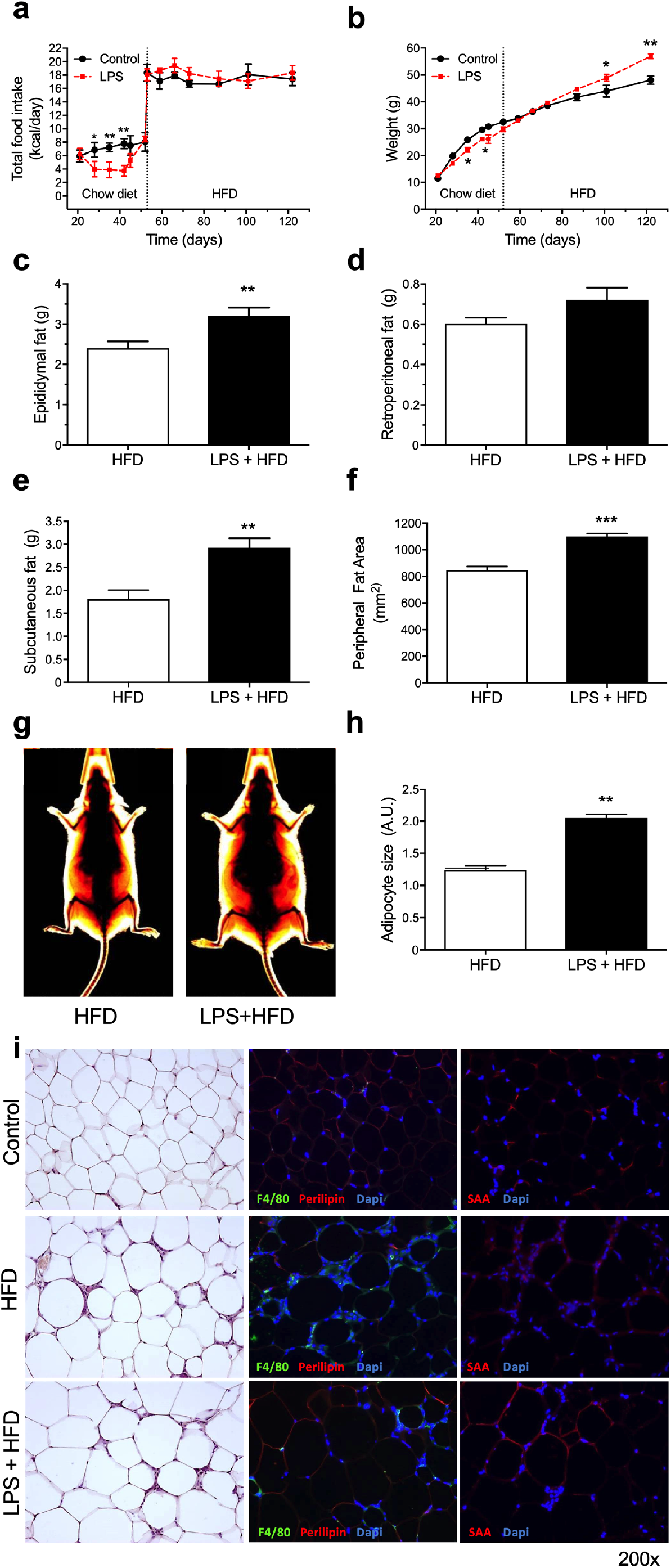
A previous history of acute endotoxaemia potentiates weight gain induced by a HFD. Mice Swiss Webster were submitted to i.p. administration of 8 consecutive doses of 10 mg/kg LPS, every 3 days, followed by 10 weeks in HFD. **(a)** Daily caloric intake, including the diet switch after acute endotoxaemia period. (b) Weight gain curve of HFD and LPS+HFD groups. **(c)** Epididymal, **(d)** Retroperitoneal and **(e)** Subcutaneous fat pad weight after HFD period. **(f)** subcutaneous fat area quantification in HFD and LPS+HFD mice after HFD period. **(g)** Representative fat area quantification. **(h)** Adipocyte size after HFD period. **(i)** Histological sections of epididymal fat pads after HFD periods showing adipocyte morphology on haematoxylin and eosin staining, macrophage infiltration (F4/80^+^) and SAA production. Data are means ± SEM from 8 mice per group (\**p* < 0.05, \*\**p* < 0.01, \*\*\**p* < 0.001, between groups, as indicated).

Under a high-fat diet, the endotoxin and SAA concentration reach serum levels about twice from that observed in lean mice (Fig. 3a compared to 1a and Fig. 3b compared to 1b) without differences between HFD and LPS+HFD groups. In a similar manner, transcript levels of *Saa3* increases upon HFD induction regardless previous administration of LPS (Fig. 3d compared to 1h). The adipose tissue expression of *Saa1.1/Saa2.1* was similar among all conditions (Fig. 3C compared to Fig. 1g). Nevertheless, mice previously submitted to multiple LPS challenges showed an increment in *Tlr-4* (Fig. 3f) and *Cd14* (Fig. 3g) mRNA expression in the adipose tissue, with no change in *Tlr-2* transcript levels (Fig. 3e). Also, mice previously submitted to multiple acute endotoxaemia had a worsened metabolic profile after the DIO protocol, with an increase in leptin (Fig. 3h) and insulin (Fig. 3l) levels, with impaired glucose homeostasis as observed by the notably affected glucose and insulin tolerance tests (Fig. 3m and *3n)*. Adiponectin (Fig. 3i), IGF-1 (Fig. 3j) and fasting glucose (Fig. 3k) concentrations were also measured in serum and no significant difference were observed.

**Figure 3.**
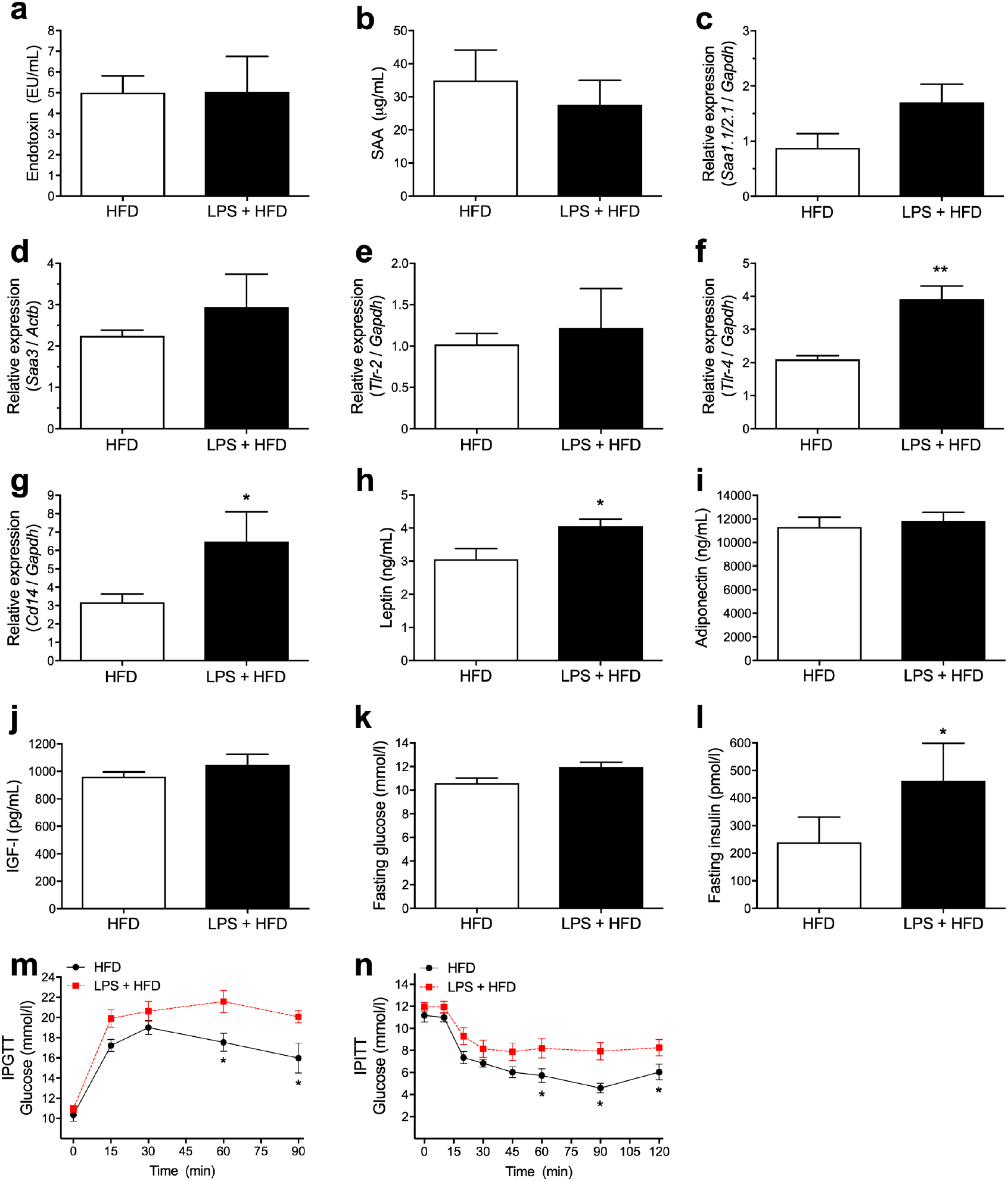
A previous history of acute endotoxaemia potentiates glucose tolerance and insulin resistance induced by a HFD. After the HFD period, mice previously submitted to multiple acute endotoxaemia were evaluated regarding its metabolic parameters. Determination of **(a)** endotoxin and **(b)** SAA in serum. **(c-g)** Quantitative Real-Time PCR for mRNA expression of **(c)** *Saa1.1/2.1* **(d)** *Saa3*, **(e)** *Tlr-2*, **(f)** *Tlr-4* and **(g)** *Cd14* in adipose tissue. **(h-l)** At last, the measurement of **(h)** leptin, **(i)** adiponectin, **(j)** IGF-I, **(k)** fasting glucose and **(l)** insulin in serum. **(m)** i.p. glucose tolerance test (IPGTT) and **(n)** i.p. insulin tolerance test (IPITT). Data are means ± SEM from 8 mice per group (\**p* < 0.05, \*\**p* < 0.01, between groups, as indicated).

### 3.3 Recurrent infection upregulates proliferative and inflammatory genes in adipose tissue

We looked at the GEO database for studies similar to our experimental protocol and where it was performed transcriptome analysis in mice adipose tissue. From the study GSE50647 (8), where mice were infected with gram-negative bacteria (*A. actinomycetemcomitans* or coinfected with *A. actinomycetemcomitans* and *C. pneumonia*), it was observed that a group of genes responsible for driving proliferation and inflammation were upregulated after infection, as well as SAA-related genes (SAA isoforms and receptors). On the other hand, the cluster of genes involved in adipogenesis was downregulated (Fig. 4).

**Figure 4.**
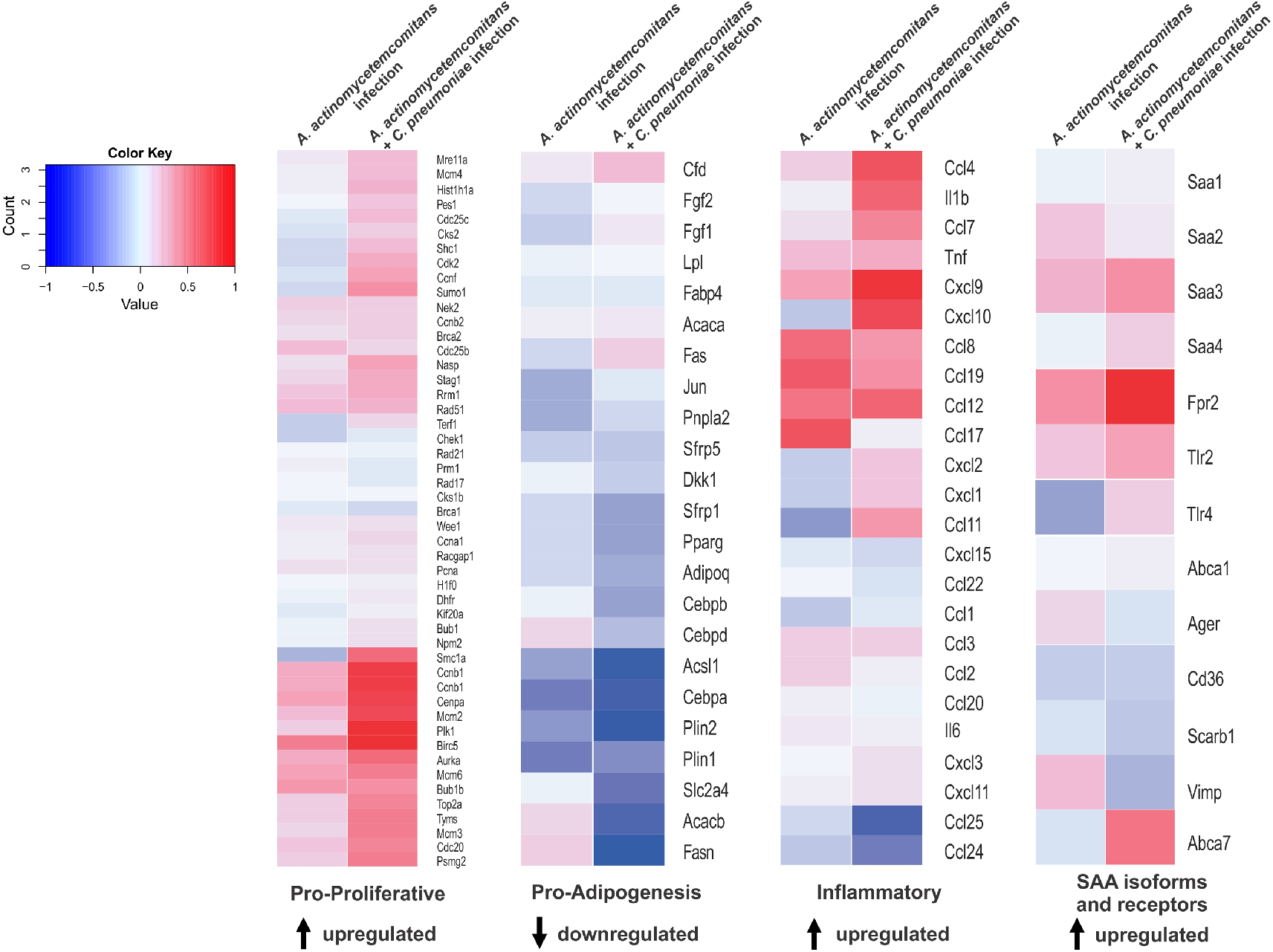
Recurrent infection modulates proliferative, adipogenic, inflammatory and SAA-related genes in adipose tissue. Gene Set Enrichment Analysis (GSEA) revealed that proliferative, adipogenic, inflammatory and SAA-related gene sets in mouse adipose tissue were significantly associated (nominal *p*-value < 0.05) with infection with *A. actinomycetemcomitans* or co-infection with *A. actinomycetemcomitans* and *C. pneumonia* (see methods for details). Heat maps show the mean log_2_ fold-change of all genes of each gene set on each condition compared to uninfected mice.

## 4. Discussion

Here we report that, besides the concurrent effects, multiple and intense endotoxaemia also causes long lasting biochemical alterations in the adipose tissue that may intensify the harmful effects of a subsequent introduced high-fat diet. Adipose tissue expression of *Tlr-4, Cd14* and *Saa3* were increased in mice submitted to multiple and severe endotoxaemia and persisted even after the end of LPS challenges. Accordingly, acute endotoxaemia, and consequently acute inflammation, should be recognized as an aggravating factor for weight gain and insulin resistance derived from a fat-enriched diet.

The intense and transient endotoxaemia detected in our experimental protocol, led to an approximately 150-times increment in endotoxin serum levels (reaching values near to 300 EU/mL) is comparable to levels found in mice and humans during infectious processes and other diseases (9). Endotoxaemia induces the release of a large amount of inflammatory mediators, such as proinflammatory cytokines and highly reactive oxygen and nitrogen intermediates identified as contributors to the LPS-induced tissue damage (10,11). The plasma level of the acute phase protein SAA increased near 1000 times and returned to basal levels after 72 hours in each LPS dose (Supplementary Table 1). During the endotoxemic phase the food intake dramatically dropped and a perceptible and expected depletion in fat depots and appearance of smaller adipocytes occurred (Fig. 1e and 3f). Besides SAA, other inflammatory markers raised in adipose tissue after the serial LPS challenges, including macrophage infiltration and *Tlr-4* and *Cd14* mRNA expression. The rapid recovery of weight and maintenance of the growth curve after the suspension of endotoxaemia tells us that the modifications derived from the acute phase, by itself, did not compromise the adipose tissue homeostasis. However, when we introduced a high-fat diet the harmful metabolic repercussions were clearly more pronounced.

TLR-4 and its co-receptor CD14 are associated with diet-induced obesity and insulin resistance in mice (1,12,13). The observed persistent increment in *Tlr-4* and *Cd14* expression in adipose tissue after LPS treatment may be one of the keys to explain the more severe weight gain and glucose homeostasis impairment. It is known that LPS and nutritional fatty acids activate TLR-4 and the co-receptor CD14 triggering the secretion of proinflammatory cytokines (14–16). This is probably one of the elements in the inflammatory signalling cascade in adipose tissue linked to metabolic diseases. Thus, the simple fact that the levels of *Tlr-4* and *Cd14* mRNA remains elevated after the interruption of acute endotoxaemia could support the higher adipose tissue responsiveness to a HFD.

Also, it is known that TLR-2 deficiency protects mice from insuin resistance and reduces tissue inflammation induced by a high-fat diet, in a process dependent of the microbiota (17). Although the expression of TLR-2 was not changed in our model of acute endotoxaemia, it is conceivable to speculate that the interaction between TLR-2 and TLR-4 might be affected, modulating its receptor cooperation and further cellular response to LPS (18).

The mechanism by which acute endotoxaemia leads to more severe consequences arising from a HFD is undoubtedly complex and may encompasses a diversity of factors. Possible endogenous TLR-4 and TLR-2 ligands involved in the signalling of acute inflammation and associate to the control of body weight may also play a role in this process. For instance, SAA is a TLR2 and TLR4 agonist (19,20) and its serum levels are positively correlated with the obesity grade (21). Moreover, although SAA is predominantly synthesized by the liver in acute inflammation, adipocytes are known producers of SAA and the mRNA and protein levels are modulated under hypoxic conditions (22,23), a common event during adipose tissue expansion. SAA production is associated with a pro-inflammatory response and also with promotion of proliferation in different cell types (5,20,24–32). Besides that, data from SAA KO mice (32,33) and studies using SAA antisense oligonucleotide (ASO_SAA_) (6,34) strongly support that SAA is part of the LPS signalling linking inflammation to obesity and insulin resistance.

In order to evaluate the comprehensiveness of some of our conclusions, especially that related to the hypothesis that acute inflammation induces preadipocyte proliferation while triggering SAA production, we performed a Gene Set Enrichment Analysis in a publicly available microarray data (GSE50647) (8) from a study in which mice were infected with gram-negative bacteria. We identified that different clusters of genes responsible for driving proliferation, inflammation and SAA-related genes (SAA isoforms and receptors) were upregulated, while genes involved in adipogenesis were downregulated after infection. Thus, the cell proliferation that occurs in adipose tissue during an infection process may provide the right condition for future susceptibility to a DIO protocol.

Considering that epidemiological data shows that low-income children have a higher prevalence of infectious diseases and are more susceptible to obesity (35,36), is inevitable to consider that excess body weight in adults may be associated with an inflammatory state during childhood. Although obesity is to a large extend a lifestyle disease, the current scientific literature has shown previous unsuspected factors contributing to obesity development. For instance, viral infections have been linked to obesity, particularly by the human adenovirus-36 (37). In humans, anti-Ad-36 antibodies are more prevalent in obese subjects (30%) than in non-obese (11%) (38). Despite differences between viral and bacterial infections it is legitimate to assume that both types of infection share some major signalling pathways linking them to obesity, such as the upregulation of TLR-4 and SAA (39,40).

In conclusion, our data describes that conditions leading to inflammation may resound for a long time in mice. If it is confirmed in humans, infections may contribute to obesity and type 2 diabetes epidemics when associated with a western diet. Furthermore, our results indicate that a more severe dietary recommendation for patients in post-infection recovery should be considered.

## Abbreviation list

(CD14): cluster of differentiation 14
(GTT): glucose tolerance test
(HFD): high-fat diet
(ITT): insulin tolerance test
(LPS): lipopolysaccharide
(SAA): serum amyloid A
(TLR-2): toll-like receptor 2
(TLR-4): toll-like receptor 4

## Acknowledgements

The authors thank the technical support from Ed Wilson Cavalcante Oliveira Santos of the University of São Paulo.

## Funding

This study was supported by Fundação de Amparo à Pesquisa do Estado de São Paulo (FAPESP) (grant number 2011/24052-4 and 2010/18498-7 [doctoral scholarship]), Coordenação de Aperfeiçoamento de Pessoal de Nível Superior (CAPES) and Conselho Nacional de Desenvolvimento Científico e Tecnológico (CNPq) (grant number 47510/2010-6).

## Duality of Interest

No conflicts of interest, financial or otherwise, are declared by the authors.

## Contribution Statement

E.M.O. contributed to the study conception and design, performing experiments, acquisition and interpretation of data and the manuscript writing. J.C.S., T.P.A, S.S., A.F.M and S.M. performed some experiments. H.T.I.N. design experiments and interpreted data. R.A.F. designed experiments and A.C. contributed to the study conception and design, interpretation of data and the manuscript writing.

