## supplementary for "Acute inflammation is a predisposing factor for weight gain and insulin resistance"

**Supplementary Figure 1A.**

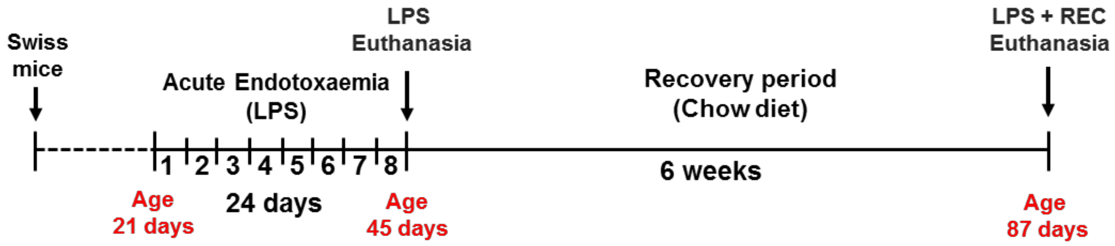


**(LPS + REC) Acute endotoxaemia followed by recovery period under chow diet.**. The method of multiple inductions of acute endotoxaemia comprises intraperitoneal administration of 8 consecutive injections (every 3 days) of LPS 10 mg/kg (Lipopolysaccharides from *Escherichia coli* 026:B6, Sigma-Aldrich®, St. Louis, MO, USA), in saline (NaCl 0,9%), starting at weaning (21 days of age) with end at 45 days of age of the animal, followed by a recovery period of 6 weeks under chow diet. For acute endotoxaemia experiments, mice were randomly assigned into 2 different groups: the Control group and the LPS group, with euthanasia occurring after the last acute phase period or after 6 weeks from the last acute phase induction (recovery period).

**Supplementary Figure 1B.**

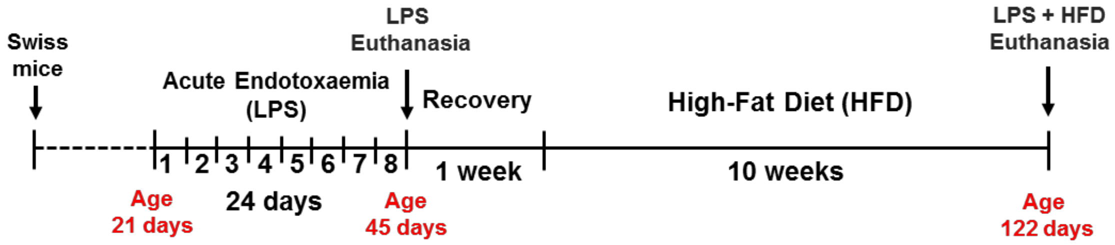


**(LPS + HFD) Acute endotoxaemia followed by High-Fat Diet (HFD).** For acute endotoxaemia followed by 10 weeks on a high-fat diet (LPS+HFD) experiments, the animals were randomly assigned into 2 different groups: HFD group and LPS+HFD group. The HFD mice were submitted to a HFD for 10 weeks starting concurrently with the LPS+HFD group. The LPS+HFD mice were underwent to multiple inductions of acute endotoxaemia followed by 1 week of recovery period in standard chow diet plus 10 weeks on a HFD.

### Supplementary Figure 2

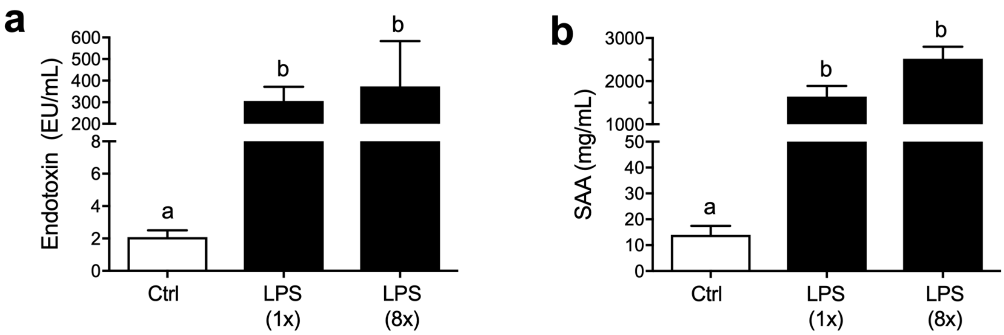

**Supplementary Table 1-** SAA profile during acute endotoxaemia

| | SAA (µg/mL) | |
|---|---|---|
| Time (hours) | Control group | LPS group |
| 0 h | 16.8 ± 8.2 | 15.4 ± 6.8 |
| 6 h | 21.7 ± 6.8 | 986.4 ± 75.8*** |
| 12 h | 25.4 ± 13.5 | 1527.0 ± 193.6*** |
| 24 h | 18.2 ± 10.7 | 888.4 ± 141.0** |
| 48 h | 7.1 ± 2.3 | 125.4 ± 40.1** |
| 72 h | 17.5 ± 8.7 | 29.6 ± 11.7 |

Data are means ± SD from 3 mice per group (***p*<0.01, ****p*<0.001, between groups, as indicated)

**Supplementary Table 1. SAA profile during acute endotoxaemia.** SAA was quantified in serum after 6, 12, 24, 48 and 72 hours of LPS-treatment (10 mg/kg).

**Supplementary Table 2-** Experimental diet composition

| Ingredients (g/Kg) | Chow diet[a] 16.7 kJ/g | High-fat diet 23.2 kJ/g |
|---|---|---|
| Sucrose | 100 | 133.56 |
| Casein | 120 | 186.98 |
| Corn oil | 80 | 53.42 |
| Lard | - | 300 |
| Cellulose | 50 | 66.78 |
| Mineral Mix (Rhoster®) | 35 | 46.74 |
| Vitamin Mix (Rhoster®) | 10 | 13.36 |
| DL-Methionine[b] | 1.8 | 2.4 |
| Choline Bitartrate | 2.5 | 3.34 |
| Tert-butylhydroquinone | 0.01 | 0.04 |
| Corn starch | 600.69 | 193.38 |

[a]According to AIN-93M.
[b]2-amino-4-methylsulfanylbutanoic acid.

**Supplementary Table 3 -** PCR primers used in all quantitative PCR assays

| Primer (gene / protein) | Forward | Reverse |
|---|---|---|
| *Saa1.1/2.1* (SAA1 / SAA2) | 5'-AGA CAA ATA CTT CCA TGC TCG G-3' | 5'-CAT CAC TGA TTT TCT CAG CAG C-3' |
| *Tlr2* (TLR-2) | 5'-CAG CTG GAG AAC TCT GAC CC-3' | 5'-CAA AGA GCC TGA AGT GGG AG-3' |
| *Tlr4* (TLR-4) | 5'-TCA TGG CAC TGT TCT TCT CCT-3' | 5'-CAT CAG GGA CTT TGC TGA GTT-3' |
| *Cd14* (CD14) | 5'-GCG AGC TAG ACG AGG AAA GT-3' | 5'-CAC GCT TTA GAA GGT ATT CCA G-3' |
| *Gapdh* (GAPDH) | 5'-TGG CAA AGT GGA GAT TGT TGC C-3' | 5'-AAG ATG GTG ATG GGC TTC CCG-3' |
